# NRF2 Regulates Contractile Vascular Smooth Muscle Cells Phenotypic Switching In Abdominal Aortic Aneurysm

**DOI:** 10.1101/2023.08.25.554907

**Authors:** Xiaoyong Xiao, Xiaoran Huang, Feier Song, Guona Chen, Yuelin Zhang, Xincheng Liu, Xueke Zhou, Yimei Hong, Haiwei He, Jinxiu Meng, Xiaojia Huang, Lintao Zhong, Xin Li

## Abstract

**OBJECTIVE:** Evidence from previous studies has demonstrated that NRF2 can protect cardiovascular progression,which prompted us to study if NRF2 is involved in AAA development.

**APPROACH AND RESULTS:** The difference and function of NRF2 in AAA were verified on human aortic tissues combined with scRNA-seq samples.We then used angiotensin II (Ang II) infusion mouse model with VSMCs-specific knockout to study the role of NRF2 in AAA formation.Primary cultured VSMCs were used to study how NRF2 regulates contractile VSMCs phenotype.Patient specimens were obtained to investigate the relevance of NRF2 expression to human AAA disease. NRF2 was induced in abdominal aortic VSMCs in both mouse and human AAA tissues. VSMCs-specific NRF2 knockout increased experimental AAA formation. Mechanistically, NRF2 regulates the contractile VSMCs phenotypic through miRNA145, leading to increased classic contractile protein level to prevent the development of AAA.

**CONCLUSIONS:** NRF2 is a negative regulator of AAA development, and thus may represent a potentially new therapeutic target to inhibit AAA growth and rupture.

---

Abdominal aortic aneurysm (AAA) is a permanent and localized dilation of the abdominal aorta that can lead to a fatal rupture of the aorta^1,2^. At present, the treatment of AAA mainly relies on surgery^3^. Due to the lack of effective drugs to treat AAA, patients still face problems such as aortic enlargement and aneurysm recurrence after surgery, which seriously affect the prognosis of patients.

The pathological mechanism of AAA is complex, and its basic pathological features include remodeling of vascular matrix, vascular inflammation and functional changes of vascular smooth muscle cells (VSMCs)^4–7^. VSMCs is the main cellular components of the aortic media. Current studies suggest that the functional changes and phenotypic transition of VSMCs are the most important mechanisms of the occurrence and development in AAA^8–13^. Although studies have confirmed that VSMCs lesions are involved in the pathological process of AAA, in-depth studies on the potential gene regulation of VSMCs function at the specific cellular level are still lacking.

Nuclear factor E2-related factor 2 (NRF2) is a key transcription factor that coordinates the basal and stress-induced activation of a large number of cell protection genes^14–16^.Under normal circumstances, NRF2 is located in the cytoplasm and will be ubiquitinated by BCR (Keap1) complex and degraded in the cytoplasm, so the level of NRF2 protein is low in many types of cells^17,18^.Studies have showed NRF2 is an important component of the defense against cardiovascular diseases, such as atherosclerosis, hypertension,intracranial aneurysm and heart failure^19–24^. However, there is study reveal that high NRF2 expression activation does not always lead to positive outcomes and may accelerate the development of certain neoplastic diseases and cardiovascular diseases (such as atherosclerosis)^25,26^. Just as NRF2 can protect normal cells from damage, it can also prevent pathogenicity cells from injure. When cells undergo switching, they are exposed to many stressors, and abnormal upregulation of NRF2 can promote the switching, which can help disease-causing cells grow, proliferate, and resist treatment. Previous studies have found that NRF2 can regulate the function of VSMCs,which inhibit the proliferation, migration,calcification,aging and apoptosis in VSMCs. Recent studies have found that NRF2 deficiency can increase the risk of AAA occurrence and rupture, but the specific role and mechanism of NRF2 in AAA are not fully understood^27–29^.

We first verified the differences and function of NRF2 at the cellular level by Single-cell RNA sequencing(scRNA-seq).ScRNA-seq technology refers to the sequencing of single-cell genome or transcriptome to obtain genome, transcriptome or other multi-omics information to reveal cell population differences and cell development lineage relationships^30–32^. At present, scRNA-seq is widely used in large vascular diseases, which can not only detect the heterogeneity among single cells, identify rare cells, map cells, but also help to discover potential biomarkers and drug targets^33–35^. In addition, in the study of aortic aneurysm dissection, it was found that scRNA-seq technology can also analyze the cell type expressing specific genes as well as its functional status and mechanism of action, so as to provide more accurate therapeutic targets and scientific basis for preventing the occurrence and development of aortic aneurysm dissection^36–38^.

This study analyzed the characteristics of the whole aortic wall subpopulation cells from scRNA-seq data, further revealed the functional characteristics and possible targets of VSMCs with high NRF2 expression in AAA group, and found clues that NRF2 may regulate the phenotypic switching of VSMCs to promote the progression of AAA, which was verified experimentally.

## Materials and Methods

### Human specimens

Human healthy abdominal aorta and AAA specimens were obtained from surgical operations of patients with abdominal aortic aneurysms in the Department of Surgery, Guangdong Provincial People’s Hospital (Guangdong Academy of Medical Sciences), Southern Medical University. The control samples were aortic tissue of donors.All included patients underwent surgical resection at Guangdong Provincial People’s Hospital, GuangZhou,China, between January and May 2021.All participants gave written informed consent before the specimens were collected. All specimens were collected under a protocol approved by the Ethics Committee of Guangdong Provincial People’s Hospital (No.KY-Z-20210219-02).Aorta proteins were extracted from formalin-fixed tissues by following the published protocol.

### Tissue dissociation and preparation of single-cell suspension

3 AAA and 3 control(2 donors and 1 recipients)samples obtained were used for single-cell analysis.The obtained tissue was cut into vascular rings under a stereoscope. Then, the vascular rings were transferred into EP tubes and were washed with DPBS. For tissue dissociation^39^, Pierce™ Primary Cardiomyocyte Isolation Kit (Thermo Fisher Scientifific™, Cat. No. 88281) was used following the manufacturer’s instructions. Afterward, we performed blind shear and bathed it at 37 °C for 25 min. After the digestion process, 400–500 μL DPBS was added to the mixture, pipet-mixed gently, and then fifiltered with a 40-μm fifilter to collect the dissociated cells. Thereafter, the mixture was centrifuged for 5 min (4 °C, 400 × g). The supernatant was discarded and a 200 μL cold Sample Buffer (BD R added hapsody™, Cat. No. 650000062) was added to the cell mass. Finally, the mixture was pipetmixed gently to obtain the cell suspension for single-cell experiments.

### Cell capture, scRNA-seq library preparation, and sequencing

For cell capture and library preparation for scRNA-seq, the BD Rhapsody™ system (BD Biosciences, USA) ^40^was applied based on the manufacturer’s protocols. In brief, 1 μL Calcein AM (2 mM; Thermo Fisher Scientifific, USA, Cat. No. C1430) and 1 μL DRAQ7 (0.3 mM; Thermo Fisher Scientifific, USA, Cat. No. 564904) were added to 200 μL cell suspension (1:200 dilution). Then, the mixture was pipet-mixed gently and incubated at 37 °C in dark for 5 min. The cell viability and concentration of the suspension were measured by Hemocytometer Adapter (Cat. No. 633703). The single-cell suspension was then loaded onto a BD Rhapsody cartridge (Cat. No. 400000847) with >200,000 microwells. Thereafter, cell capture beads were loaded into the microwells and washed to make sure that one magnetic bead binds to only one cell in a single microwell. Subsequently, the lysis mix was added and incubated at room temperature (15–25 °C) for 2 min. The cell capture beads were eventually retrieved for subsequent complementary DNA synthesis, exonuclease I digestion, and multiplex PCR-based library construction. Sequencing libraries were prepared using Whole Transcriptome Analysis^41^ Index PCR and the PCR product was purifified to enrich the 3′ end of the transcripts linked with the cell label and molecular indices. Finally, quality checks of the indexed libraries were performed with a Qubit Fluorometer using the Qubit dsDNA HS Assay. The sequencing of the libraries was conducted on an Illumina NovaSeq 6000 system.

### scRNA-seq data preprocessing and analysing

The raw sequencing reads were trimmed to keep the fifirst 75 bases. Then the trimmed reads were quality fifiltered using fastp. The BD Rhapsody whole tran scriptome offificial analysis pipeline (Version 1.5.1) was applied under the default settings to obtain a cell-gene expression matrix and a quality control report for each sample. The human reference genome^42^ was used for read alignment. The expression matrix was imported into Seurat^43^ for further preprocessing. We fifiltered out genes with counts in fewer than three cells to eliminate genes that were most likely discovered due to random noise. We fifiltered cell outliers (<fifirst quartile-1.5 interquartile range or >third quartile+1.5 inter quartile range) based on the proportion of mitochondrial genes, the number of expressed genes, and the sum of unique molecular identififier (UMI) counts. To eliminate doublets, cells were fifiltered out using Scrublet. Furthermore, cells enriched in hemoglobin gene expression were also excluded. We normalized the UMI count sum of each cell to 10,000 and then log-transformed the data. To identify shared cellular states across samples, all the datasets were integrated via canonical correlation analysis (CCA) implemented in Seurat. The G2M phase score, S-phase score, UMI count, and mitochondrial gene proportion were regressed using linear models.Principal component analysis (PCA) was used to reduce the dimensionality of scRNA-seq expression data. A neighborhood graph of the cells was generated based on the fifirst 30 PCA components. Using UMAP, we embedded the neighborhood graph in two-dimensional space. Louvain clustering was used to cluster the cells (resolution = 1).Using the likelihood-ratio test implemented in the function “FindMarkers” of Seurat, we detected differentially expressed genes between two groups of cells. The signifificance threshold was set to a log2-fold change >0.25 and adjusted P value < 0.05. GO was used to perform functional enrichment analysis on a gene list with a Bonferroni-corrected P value < 0.05.

### Animals

KO(NRF2^ΔVSMC^) and WT control (NRF2^flox/flox^)mice were originally provided by Saiye Model Biological Research Center. No differences in size and weight were detected between WT and KO mice. All animals were housed in isolation rooms at the Animal Facilities of the Institute of Southern Medical University. The Ethics Review Board of the Institute of Southern Medical University approved all animal procedures. All animal procedures were performed in accordance with the guidelines of Directive 2010/63/EU of the European Parliament on the protection of animals used for scientific purposes.

### Ang II-induced murine model of AAA

Eight-week-old WT and NRF2^ΔVSMC^ mices were infused with phosphate buffered saline (PBS) or Ang II (1000 ng/kg/min) via osmotic minipumps (AP-2004, Alzet, CA, USA) for 28 days, as described previously^12,44^. Briefly, mice were anesthetized with inhaled isoflurane (5% for induction and 2% for maintenance) and the minipumps were surgically implanted into the subcutaneous space of the mice in the back of the neck. 28 days later, mice were anesthetized using 2.0% isoflurane, and hair was removed from the abdomen by using depilatory cream (Nair; Church & Dwight Co, Inc; Princeton, NJ). Mice were then laid supine on a heated table, and warmed ultrasound transmission gel was placed on the abdomen. Aortic diameters were measured using a doppler ultrasound Vevo 1100 Imaging System (VisualSonics) with a real-time microvisualization scan head in B mode. The B-Mode is a two-dimensional ultrasound image display composed of bright dots representing the ultrasound echoes. The brightness of each dot was determined by the amplitude of the returned echo signal. The abdominal aortas were then harvested for RNA, protein, and morphological or histological analyses. AAA incidence was defined by an increase of external aorta diameter by 50% or greater as compared to aortas from saline-infused mice. For WT mice, prior to saline or Ang II infusion, the mice were pre-treated with tamoxifen (1 mg/day, i.p.) for 5 days to induce Cre activity and generate NRF2-VSMC heterozygous knockout mice.

### Cell culture

Abdominal aortic aneurysm tissue was harvested from AAA patients who underwent surgical repair. Samples of healthy human aortic tissue was collected from donors and served as the control group. The procedure was approved by the research ethics board of Guangdong Provincial People’s Hospital. Written informed consent was obtained from all study subject. Human VSMCs used in this study were prepared as described in our previous study^45^. Briefly, after cleaning adipose tissue and washing with PBS, the medial tissue was dissected from the adventitia and intima. Next, the media was cut into 1-2 mm^3^ pieces and carefully transferred to 25cm^2^ culture flask and incubated for adhesion at 37°C for 1 h. After the tissue attaching to the culture flask, the medial tissues were gently cultured with Dulbecco’s modified Eagle medium (DMEM; Gibco) supplemented with 15% fetal bovine serum (FBS; Gibco) and 100 µg/mL penicillin and streptomycin (P/S, Thermo Fisher Scientific). The medium changed approximately every 3 to 4 days. VSMCs migrated out from the medial tissues within 1-2 weeks. The pieces were then removed and the cells were regularly collected and passaged. All VSMCs at passage 2-4 were used in this study.

### Real-time PCR

Human AAA tissues, as well as aortic wall tissues from healthy human controls, were snap-frozen in liquid nitrogen; homogenates (0.2 g) were resuspended in TRIzol buffer (Life Technologies); and total RNA was purified. Similarly, lysates from VSMCs were resuspended in TRIzol,and total RNA was purified. Duplicate samples were quantified by determining absorbance at 260 nm, and real-time PCR was performed as described^46^.

### Western blotting

Abdominal aorta or VSMCs were lysed in RIPA lysis buffer (1% Nonidet P-40, 0.1% sodium dodecyl sulfate (SDS), 0.5% sodium deoxycholate, 1 mm sodium orthovanadate, and protease inhibitors) to extract total proteins. Samples were separated on SDS-polyacrylamide gels and electro-transferred onto nitrocellulose membranes (Amersham Biosciences). After blocking with 5% BSA, the membranes were incubated with a primary antibody at 4 °C overnight. The membranes were then incubated with IRDye secondary antibodies (LI-COR Biosciences) at room temperature for 1 hour. The protein expression was detected and quantified by Odyssey CLx Imaging System (LI-COR Biosciences)^47^.

### RNA-Seq

Total RNA was extracted from VSMCs of WT and KO mices after transfection with (n = 3/group) using the RNeasy MiniKit (QIAGEN, 74106) and then treated with RNase-free DNase I (QIAGEN, 79254) according to the manufacturer’s instructions. In brief, RNA quality was determined by BioAnalyzer (Agilent). Then the RNA library was prepared with the NEBNext Ultra RNA Library Prep Kit (New England Biolabs), and 51 bp read paired-end sequencing was performed on a HiSeq 6000 platform (Illumina). A total of 266 million reads were generated, with an average of 38 million reads per sample. RNA-Seq read mapping was performed as described previously^48^. In brief, FastQC (Babraham Bioinformatics) was used for quality control of the sequencing reads from each sample. Gene expression quantification was performed using Salmon^49^, with mouse cDNA sequences of GRCh38 (Ensembl database) as reference. Differential expression analysis was performed with the DeSeq2 package in R^50^. GSEA was performed to interpret gene expression profiles of WT and KO mices VSMCs. Genes were mapped to the HALLMARK and GO gene set in the MSigDB for pathway analysis^51^.

### Immunohistochemistry

Primary antibodies were anti–IL6, CD68, and MMP2. Rabbit anti-goat horseradish peroxidase (HRP), donkey anti-rat HRP (The Jackson Laboratory), and Alexa Fluor 488 donkey anti-rat (Invitrogen) were used as secondary antibodies. HRP was then added, and sections were stained with 3,3′-diaminobenzidine (DAB) substrate-chromogen (DAKO) and counterstained with hematoxylin. Computer-assisted morphometric analysis was performed with Image-Pro Plus software^52^. The threshold setting for area measurement was equal for all images. Samples were examined in a blinded manner. Results were expressed as % positive area versus total area (CD68,IL6 and MMP2).

### Statistical analysis

Results are expressed as means ± SE. For analysis of data from human samples or from the experimental models, groups were compared with the Mann-Whitney test. In vitro experiments were replicated at least three times and analyzed by Student’s *t* test or the analysis of variance (ANOVA) test followed by Tukey’s as appropriate. Statistics were performed using SPSS software (23.0; SPSS Inc., Chicago). Significance was defined as *P* < 0.05 (two-tailed).

## Result

### 1. NRF2 expression was significantly increased in VSMCs of AAA

First,we collected aortic tissue samples from patients with human abdominal aortic aneurysm and normal subjects for PCR and Western blotting in vitro. Comparative analysis showed that the expression level of NRF2 in AAA group was significantly higher than the level in the control group (Figure.1A,B). Then we performed scRNA-seq on 3 pairs of AAA and control samples from human full-layer aortas (3v3), and mined the public database dataset GSE166676 (containing 4 abdominal aortic aneurysms and 2 normal full-layer aortic wall scRNA-seq). We combined the two dataset after quality control using t-SNE plot to obtain a total of 32487 cells, including 12997 cells in AAA group and 19490 cells in control group.Subsequently,all cells were grouped into 10 groups, namely B cells (CD79A,CD37), Fibroblasts (DCN,COL1A2),SMC (MYL9,ACTA2), T Cells (CD3D,CD2), Endothelial Cells (VWF,IFI27), Macrophages Cell (LYZ,CD68), Monocytes Cells(S100A9), Plasma Cells (MZB1), Mast Cells (CPA3), Dendritic Cells (GZMB)(Figure.1C). These cells all have high expression markers as noted above(Tab1,Figure.1D,E).We further calculate the proportion of each cell between two samples showed that non-immune cells such as Fibroblasts,SMC(Smooth Muscle Cells) and EC (Endothelial Cells) decreased significantly in AAA(Tab2,Figure.1F, Sup.1A). However, immune cells such as B cells, T cells, Macrophages, Monocytes, Plasma Cells, Mast cells, and Dendritic Cells were significantly increased in AAA. At the same time, gene differential expression analysis of above cells showed that NRF2 had significant differences in VSMCs, and was significantly higher in AAA, while the differences were not obvious in other cells (Figure.1G,Sup.1B). These data suggest that NRF2 may be involved the AAA development especially in VSMCs of human patients.

**Figure 1.**
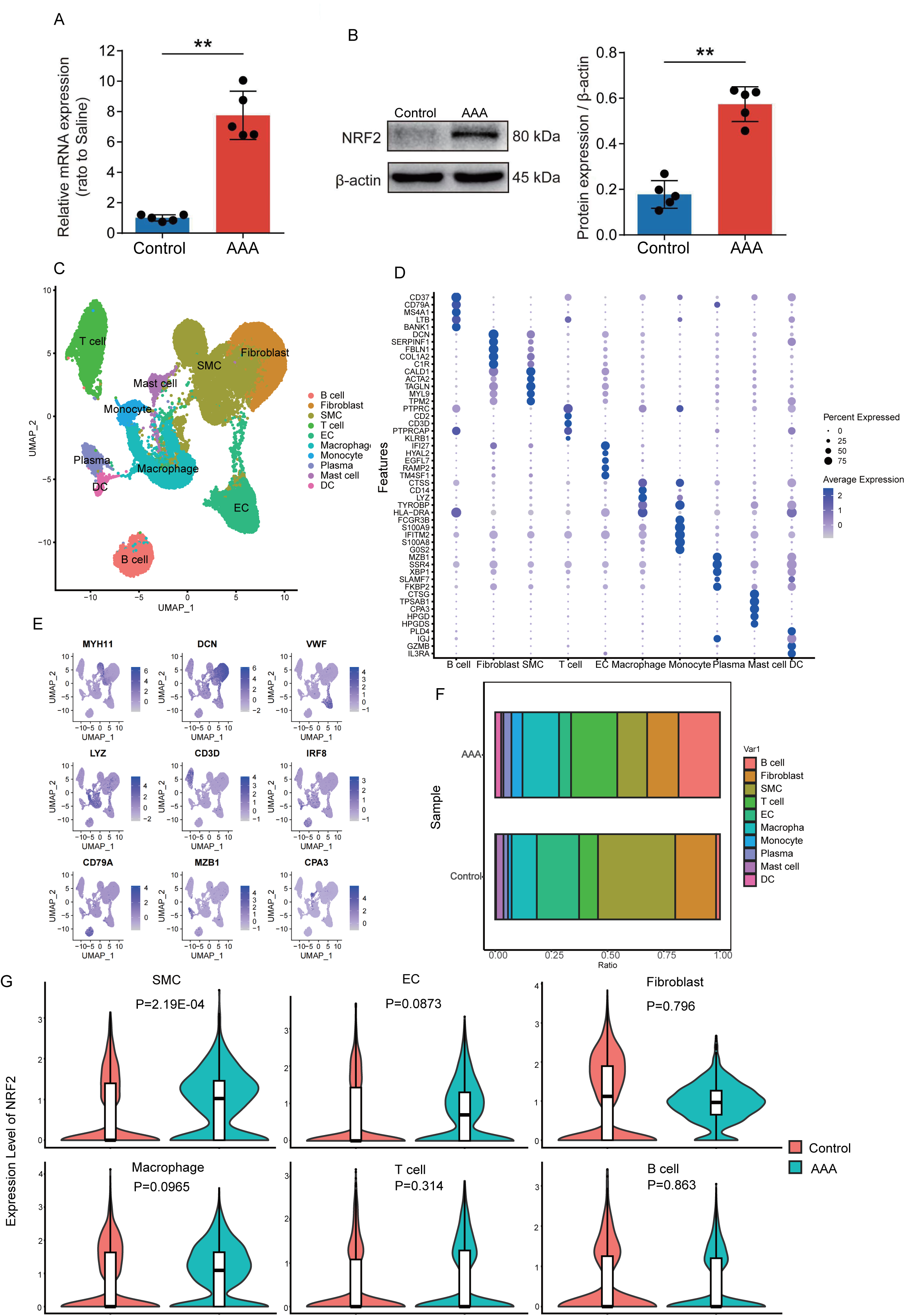
NRF2 expression was significantly increased in VSMCs of AAA. (A,B) PCR and western-blotting showing the expression levels of NRF2 in human aorta tissue. N=5, p*<0.05; student t-test. (C) A umap plot showing all cells colored according to the 10 major cell types. (D) Mean expression of selected genes in the major cell types. (E) Relative expression of several marker genes in all cells from all samples. Cells were projected onto a umap plot. (F) The composition of different cell type is shown in the horizontal bar plot. (G) The difference expression of NRF2 in each cell type between AAA and Control groups in VlnPlot.

### 2. Differential function of NRF2 in VSMCs of AAA samples

In order to explore the specific function of NRF2 in VSMCs,VSMCs were then isolated in group AAA of single-cell transcriptome sequencing.Followly,VSMCs was divided into NRF2+ group and NRF2-group according to NRF2 expression, among which NRF2+ group accounted for 36% (Figure.2A,Sup.1C). Subsequently, pseudo-time series analysis showed that NRF2+ group in AAA was in the upstream position of differentiation, while NRF2-group was in the downstream position of differentiation (Figure.2B-E). Further differential gene analysis between the two groups showed that NRF2+ group mainly expressed contractile factors such as MYH10,MYL9 and stress factors such as ATF4,ATF3, while NRF2-group mainly expressed inflammatory factors such as CCL19,CXCL13 and matrix metalloproteins such as MMP3 (Tab3,Figure.2F). GO enrichment analysis showed that NRF2+group mainly enriched signals such as smooth muscle contraction, oxidative stress, and cell differentiation, while NRF2-group mainly enriched signals such as matrix remodeling and inflammation response(Figure.2G).

**Figure 2.**
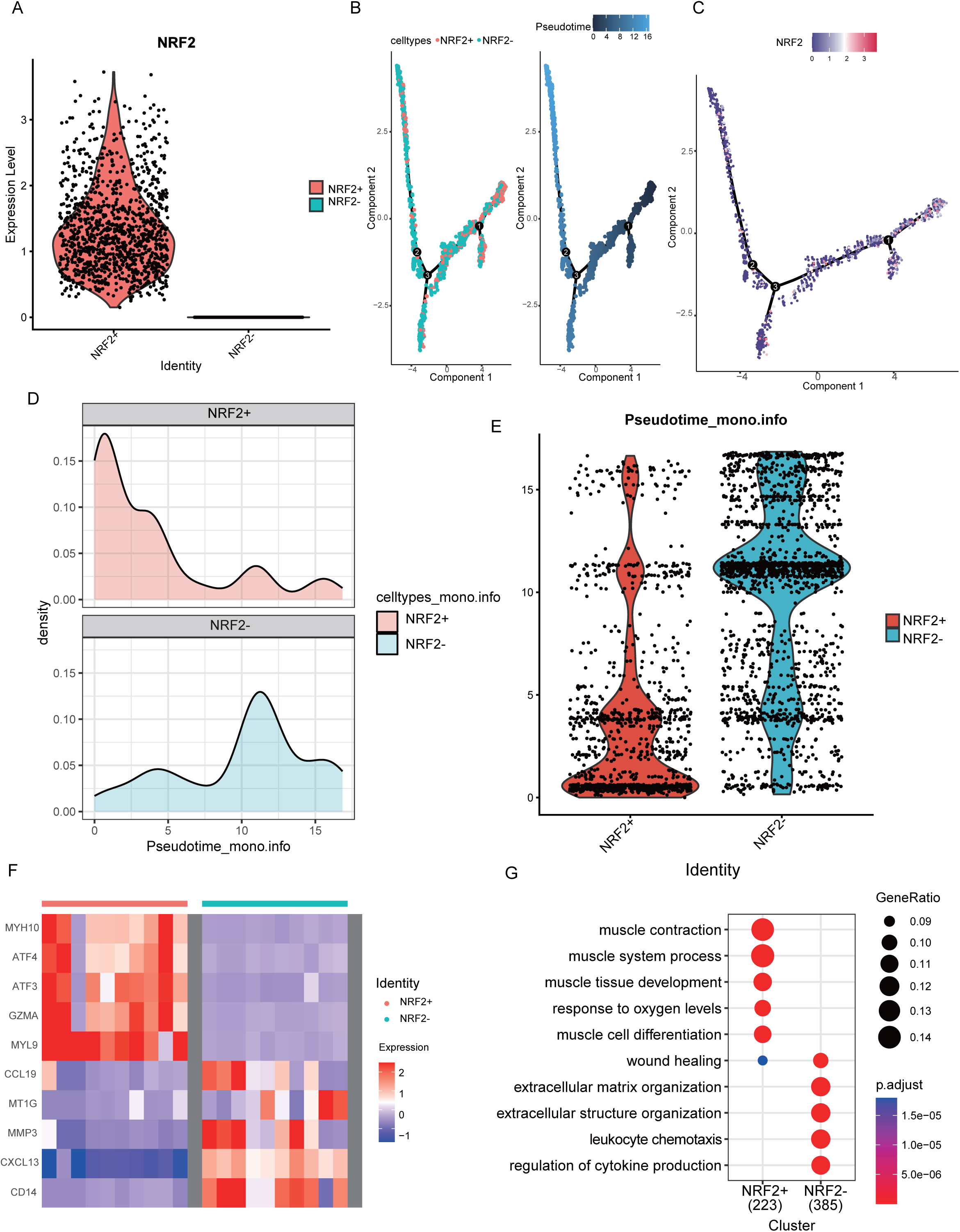
Differential function of NRF2 in VSMCs of AAA samples. (A) VSMCs was divided into NRF2+ and NRF2-group according to its expression in AAA. (B-E)Pseudo-time series analysis showed that NRF2+ group in AAA was in the upstream position of differentiation, while NRF2-group was in the downstream position of differentiation. (F) DoHeatmap of differentially expressed genes (DEGs) between NRF2+ and NRF2-group. (G) Dotplot of GO analysis between NRF2+ and NRF2-group.

### 3. NRF2 promotes the VSMCs contraction phenotype

We further subdivided VSMCs into 7 subgroups to further clarify the role of NRF2 at the cellular level.Namely, fibroblast cell-like VSMCs (F-SMCs), contractile VSMCs 1(C-SMCs1), contractile VSMCs 2(C-SMCs2), macrophage cell-like VSMCs (M-SMCs), stem cell-like VSMCs (S-SMCs),T cell-like VSMCs (T-SMCs) and plasma cell-like VSMCs (P-SMCs) (Figure.3A).The proportion of F-SMCs and C-SMCs in AAA decreased significantly,while S-SMCs,T-SMCs and P-SMCs are entirely opposed outcomes(Tab4,Sup.2A,B). The F-SMCs cluster highly expressed collagen genes such as COLA1 and FBLN1,and the main enrichment function are extracellular matrix organization(Tab5,Figure.3B,C).The C-SMCs1 cluster shared highly expressed contraction-related genes such as ACTA2, CNN1,and also expressed low levels of collagen,its major function are muscle contraction process and slight enrich extracellular matrix organization.While the C-SMCs2 cluster showed the similar level of contraction and collagen genes,and enrichment equivalent function of muscle contraction and extracellular matrix organization.Further differential gene analysis between the two groups showed that NRF2 mainly had significant differences in C-SMCs1 and C-SMCs2, but no significant differences in other cell subsets (Figure.3D). Ligand receptor analysis of these seven cell subpopulations showed a significant ligand receptor relationship between NRF2 and the classical contraction factor ACTA2 (α-SMA) in VSMCs (Figure.3E). In vitro PCR and WB experiments of aortic smooth muscle cells showed that the classical contraction factors α-SMA, CNN1 and SM22α were significantly decreased after NRF2 was knocked down,while the expressions of α-SMA, CNN1 and SM22α were increased after NRF2 was overexpressed (Figure.3F,G).

**Figure 3.**
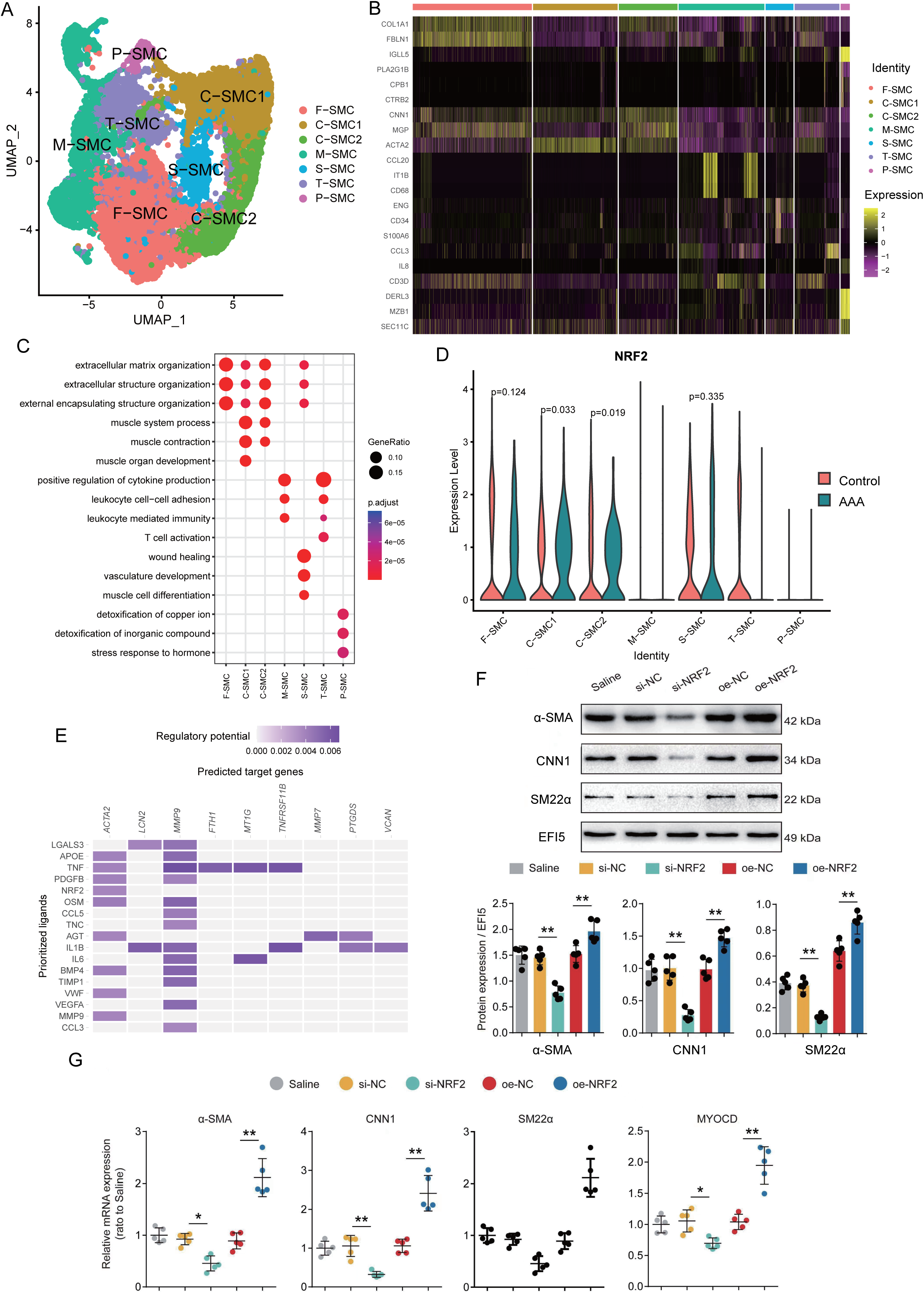
NRF2 promotes the VSMCs contraction phenotype. (A) Integrative analysis and re-clustering for VSMCs in scRNA-seq based on gene expression profile. Seven main VSMCs populations, fibroblast cell-like VSMCs (F-SMCs), contractile VSMCs 1(C-SMCs1), contractile VSMCs 2(C-SMCs2), macrophage cell-like VSMCs (M-SMCs), stem cell-like SMCs (S-SMCs),T cell-like VSMCs (T-SMCs) and plasma cell-like VSMCs (P-SMCs), were identified. (B) Mean expression of selected genes in the major cell types of VSMCs. (C) Dot plot of GO analysis in the major cell types. (D) Comparative analysis of NRF2 differential expression levels in each cell type. (E) Ligand and receptor interaction in VSMCs. (F) Western-blotting showing the protein expression levels of α-SMA, CNN1 and SM22α after NRF2 knockdown or overexpression in VSMCs.N=5, p*<0.05; student t-test. (G) PCR showing the mRNA expression levels of α-SMA, CNN1, SM22α and MYOCD after NRF2 knockdown or overexpression in VSMCs.N=5, p*<0.05; student t-test.

### 4. VSMCs-Specific NRF2 deficiency promotes AAA formation In Vivo

To determine whether the downregulation of NRF2 mediates AAA injury or acts as a self-defensive anti-AAA signal, we tested the effects of VSMCs-specific NRF2 knockout (NRF2^ΔVSMC^) on AAA formation(Sup.3A,B). We found that VSMCs-specific NRF2 knockout significantly aggravated AAA formation, suggesting that NRF2 acted as a positive regulator of AAA pathogenesis. Thus, we focused on the role of VSMCs derived NRF2 in AAA formation and development in the following experiments. In an AngII-induced AAA model, AngII infusion was administered to homozygous NRF2^ΔVSMC^ mice and their littermate controls (NRF2^flox/flox^) for 28 days, and blood pressure were monitored. AngII increased systolic blood pressure similarly in NRF2^ΔVSMC^ and NRF2^flox/flox^ mice(Figure.4A).Compared with those in NRF2^flox/flox^ mice, increased AAA prevalence and decreased survival rate were observed in NRF2^ΔVSMC^ mice(Figure.4B,C).Deteriorated AAA formation was further validated by in vivo imaging analysis.Analysis results revealed that VSMCs-specific NRF2 knockout increased maximal diameter and abdominal aortas elastic fiber degradation score in AngII-administered mouse aortas(Figure.4D-G).

**Figure 4.**
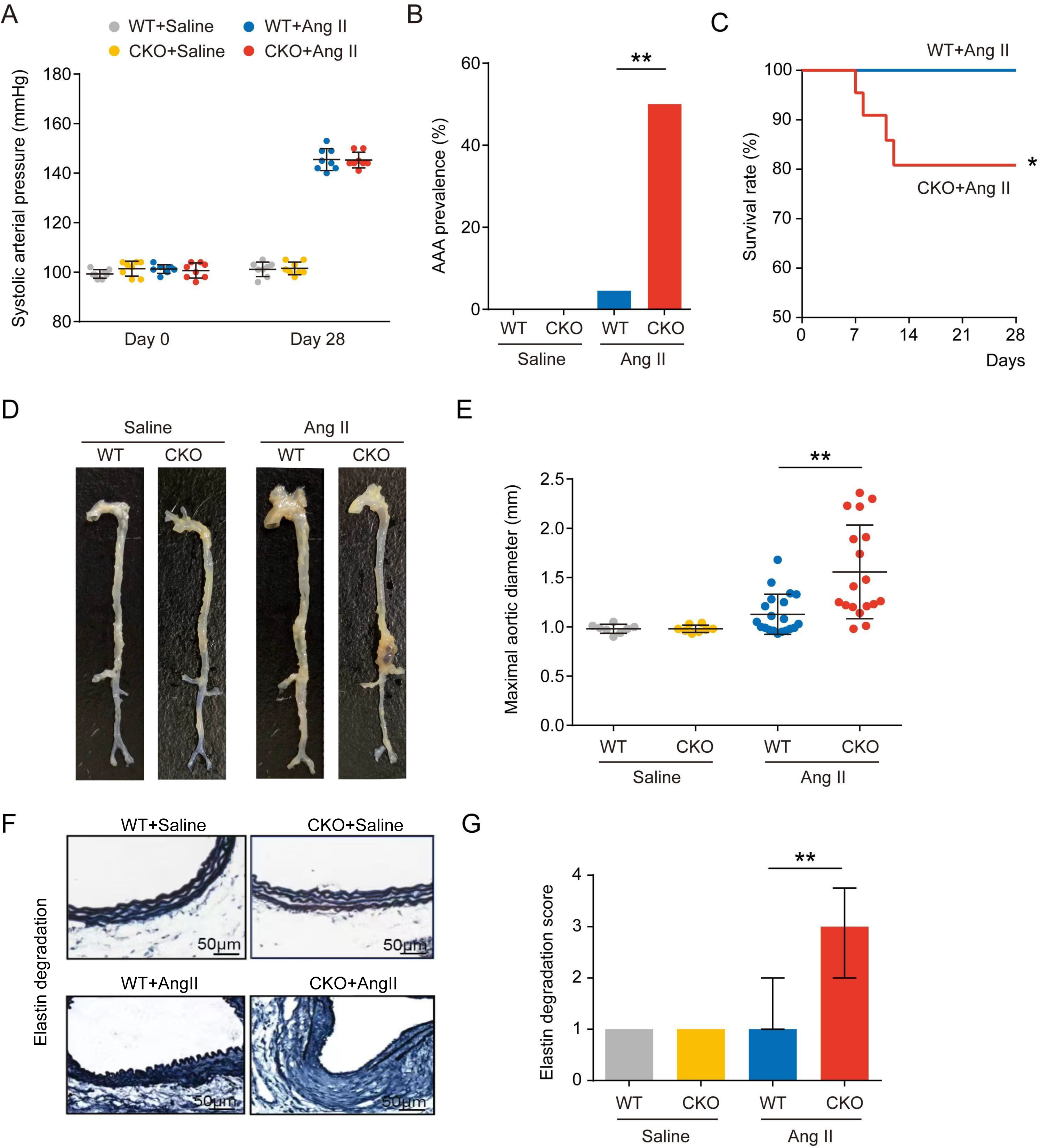
VSMCs-Specific NRF2 Deficiency Promotes AAA Formation In Vivo. (A)Abdominal aortas were collected 28 days after Ang II infusion in NRF2^ΔVSMC^ and NRF2^flox/flox^ mice.AngII increased systolic blood pressure similarly in two groups.*P >0.05. (B,C)Quantitative analyses of AAA prevalence and decreased survival rate were observed,*P = 1.18E-06,**P = 6.71E-03. (D,E)NRF2^ΔVSMC^ increased Ang II infusion-caused aorta dilation in Vivo as imaged after quantitative analyses,*P = 3.62E-02. (F,G)Abdominal aortas elastic fiber degradation scores were significantly increased in NRF2^ΔVSMC^ mouse aortas after AngII administered,*P = 1.34E-04.

### 5. VSMCs-Specific NRF2 deficiency promotes phenotypic pwitching and inflammatory response

We conducted RNA sequencing of NRF2 Deficiency in VSMCs to confirm whether AAA appear phenotypic transition after NRF2 deficiency, and found that there were a significant decrease of classic contraction factors and increase of inflammatory mediators in NRF2*^ΔVSMC^* mice(Tab6,Figure.5A,Sup.4A).GO enrichment analysis showed that the most significant downregulated function were regulation of smooth muscle contraction and smooth muscle cell differentiation,while the corresponding upregulated function were inflammatory response and muscle cell proliferation (Tab7,Figure.5B).To evaluate the VSMCs phenotypic switching in the aortic wall, we determined the expression of MYH11,α-SMA and CNN1. PCR and Western blotting results revealed that MYH11,α-SMA and CNN1 expression were dramatically abolished in the aorta of AngII-infused NRF2*^ΔVSMC^* mice(Figure.5C,D). Moreover, NRF2 deficiency promotes the expression of inflammatory factors (ie, CD68[macrophage antigen CD68], IL-6 [interleukin-6] and MMP2 [matrix metallopeptidase 2]) in the abdominal aortas by immunostaining monitored(Figure.5E).

**Figure 5.**
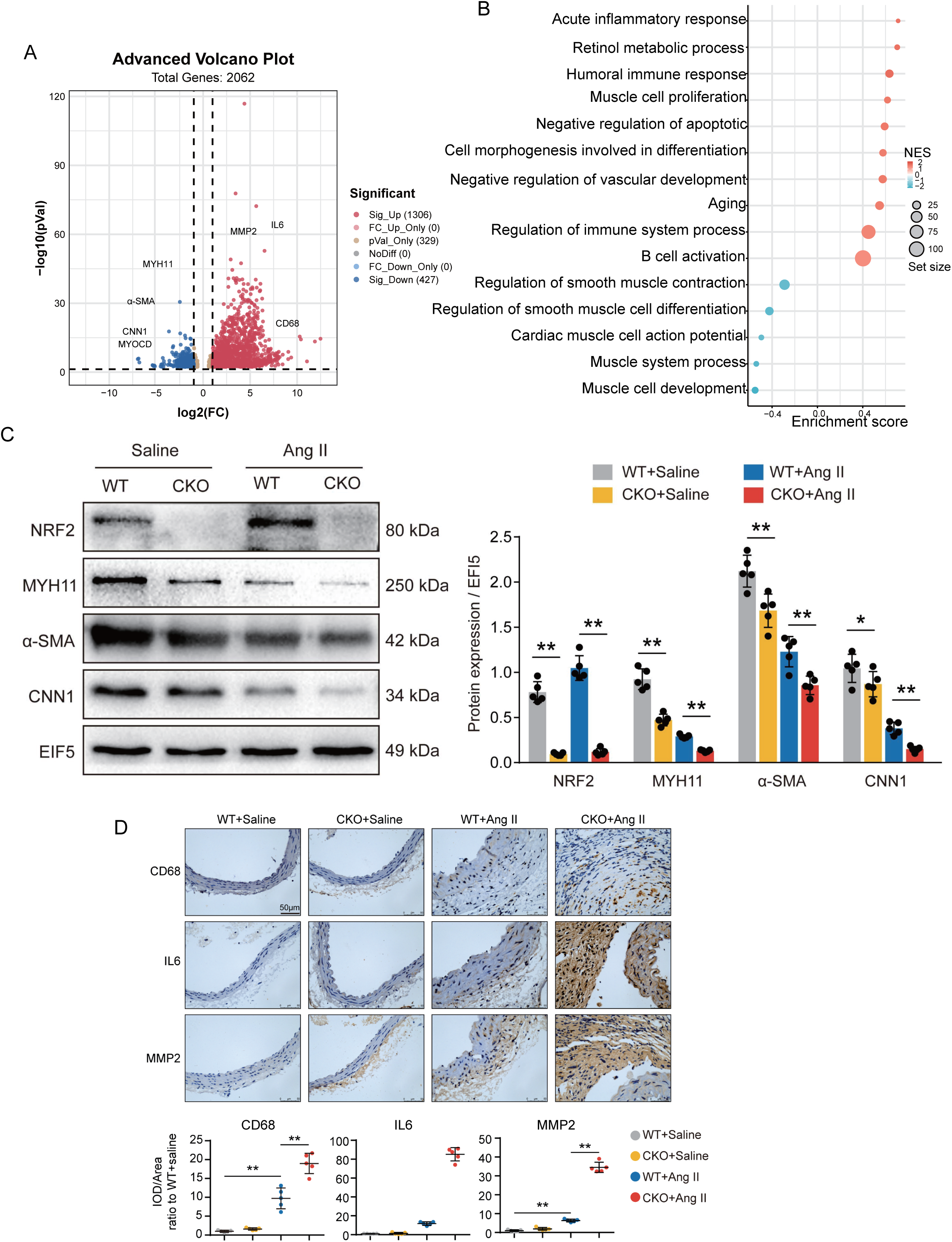
VSMCs-Specific NRF2 Deficiency Promotes phenotypic transformation and inflammatory response. (A) RNA-seq analysis of abdominal aortas from NRF2^ΔVSMC^and NRF2^flox/flox^ mice. Volcano plot showing differential gene expression between two groups. (B) Dot Plot of GO analysis in NRF2^ΔVSMC^. (C) Western blotting results revealed that MYH11,α-SMA and CNN1 expression were dramatically abolished in the aorta of AngII-infused NRF2*^ΔVSMC^* mice,*P = 5.16E-04, **P = 2.39E-06. (E)Immunohistochemical showing CD68,IL-6 and MCP-1 were dramatically increased in the aorta of AngII-administered NRF2*^ΔVSMC^*mice, *P = 3.51E-07.

### 6. NRF2 regulates the contractile VSMCs phenotypic through miRNA145

To obtain deeper insight into the underlying mechanisms linking NRF2 to the pathogenesis of AAA, bulk RNA-sequencing of mouse aortas was performed,and found 10 miRNA were significantly down-regulated in NRF2-KO group (Tab8,Figure 6A). Bioinformatics analyses of differentially expressed genes suggested that the upregulated genes were enriched in MYCOD pathways (Tab9,Figure 6B).Then,We inquire miRactDB database showed significant correlations between four miRNA and Myocd, the miRNA145 being the most correlated (Figure 6C).Previous study has shown that NRF2 has a MYOCD binding site on VSMC^72^(Sup.4B).We continued to analyze the correlation in the scRNA-seq data, showing a significant correlation between miRNA145 and NRF2 at VSMCs (Figure 6D). In vitro human cell experiments showed that si-NRF2 combined with miRNA145 inhibition resulted in a decrease in classical contraction factor expression(α-SMA,CNN1 and SM22α), while overexpression of miRNA145 resulted in an increase in classical contraction factors expression, which in a direction consistent with the expression of Myocd (Figure 6E,F).

**Figure 6.**
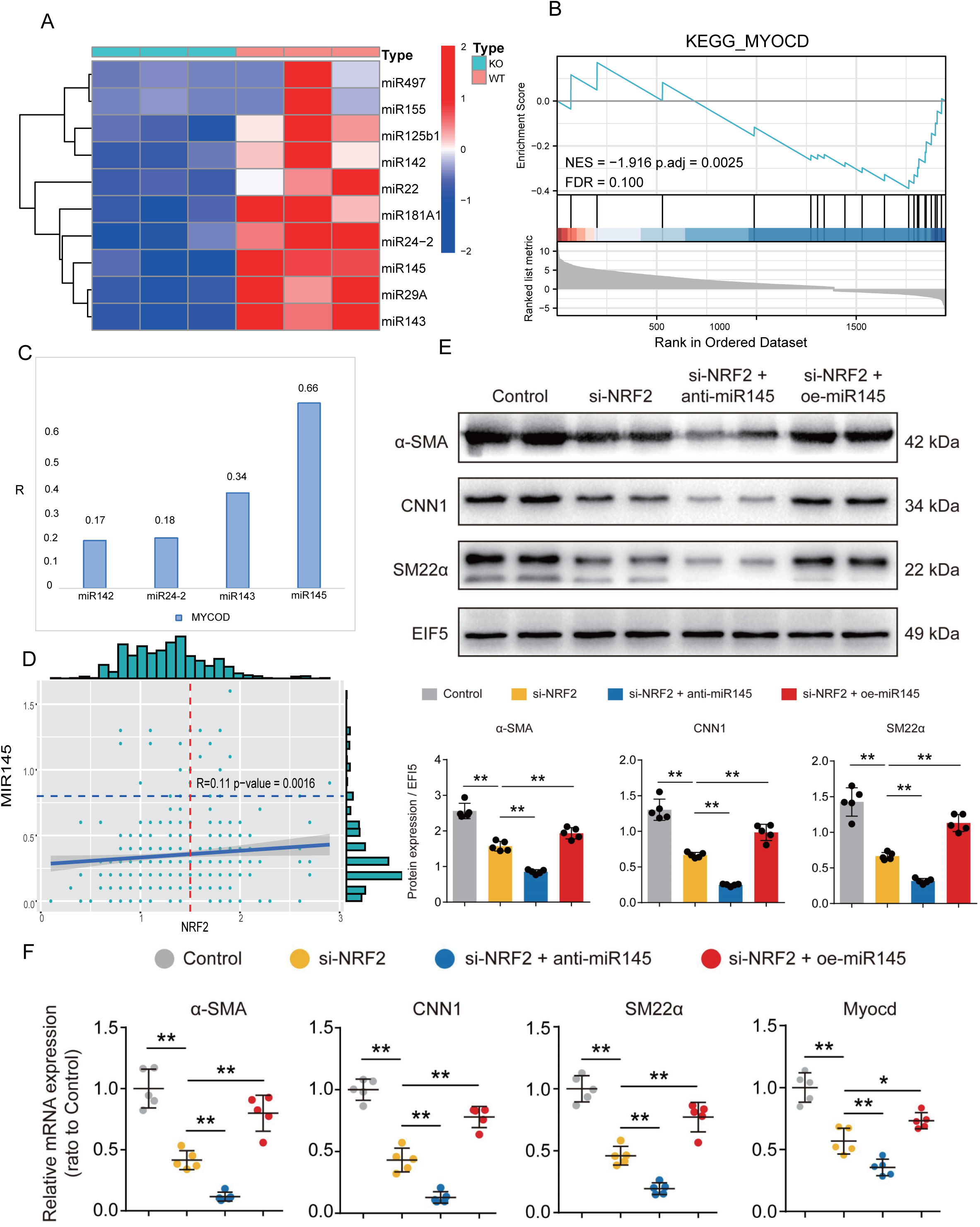
NRF2 regulates the contractile VSMCs phenotypic through miRNA14. (A)Heatmap of RNA-seq analysis revealing 10 miRNA were significantly down-regulated in NRF2-CKO group. (B) KEGG analysis of differentially expressed genes suggested that the upregulated genes were enriched in MYCOD pathways,NES =-1.916,p.adj = 0.0025,FDR = 0.100. (C)MiRactDB database showed the miRNA145 being the most correlated between four miRNA and MYOCD,R=0.66,P<0.05. (D)Sc-RNAseq data showing a significant correlation between miRNA145 and NRF2 at VSMCs,R=0.11 p−value = 0.0016. (E,F) PCR and Western blotting of vitro human cell experiments showing si-NRF2 combined with miRNA145 inhibition resulted in a decrease expression of α-SMA,CNN1 and SM22α, while overexpression of miRNA145 resulted in an increase these expression, which in a direction consistent with the expression of Myocd,N=5,p*<0.01.

## Discussion

In this work, we confirmed a novel role of the NRF2 in the pathogenesis of AAA. Novelty points include the following.First, we identified that NRF2 expression exhibited the highest fold change in human VSMCs from AAA tissues. Moreover,We confirmed the difference on scRNA-seq and verified the functional characteristics of NRF2 at the VSMCs level,which suggested that NRF2 regulates AAA through VSMCs phenotypic switching. Furthermore,VSMCs specific knockout of NRF2 promoted mice developed AAA formation in AngII-induced AAA models.Last,mechanistic studies identified miRNA145, a key miRNA that regulates phenotypic transition of VSMC mediated by MYCOD,as a downstream target trans-promoted by NRF2 that mediated the negative effects of on contractile VSMCs phenotypic and AAA formation.Collectively, the results of the our study provided the first evidence for a novel regulatory NRF2-miRNA-MYOCD signaling axis in AAA formation and progression.

VSMCs always manifested as contractile state with slight function of accretion and migration in normal human aortic tissue^53,54^.Normal VSMCs overexpression of contractile proteins can form a large number of fibroblast muscle bundles, also secrete a large number of ECM proteins to maintain the normal structure of aorta^55^. Earlier studies have confirmed that VSMCs phenotypic switching occur after VSMCs injured, which is manifested by the decline of contractile proteins, the increase of collagen synthesis, and the appearance of irregular morphology with a large number of rough endoplasmic reticulum and Golgi complex^56,57^. Studies have also shown that aorta can not adjust to normal diameter in response to blood pressure changes after the phenotypic switching of aortic VSMCs due to the loss of contractile protein, resulting in the occurrence of aorta aneurysms^56,58^. In addition, the loss of contractile protein can lead to the decrease of the density of VSMCs adhesion and cell matrix adhesion, thus contribute to the internal structure of the aorta unstable and prone to matrix remodeling. This study also confirmed that VSMCs lesions were particularly obvious in AAA through the scRNA-seq results of human aorta, which was consistent with the results of previous studies. We further isolated VSMCs for cluster analysis, showing that the proportion of contractile-SMCs and fibroblast-SMCs decreased significantly in AAA, while the proportion of inflammatory type VSMC did not change significantly. Therefore, it was convinced that the occurrence and development of AAA was mainly based on the switching of VSMC contractile phenotype.

NRF2 has been proved to protect in various progression diseases, including diabetes, tumor, trauma,immune disorders and so on^15,59–63^.In recent years, many studies have revealed the protective role of NRF2 in cardiovascular diseases, which can play an antioxidant, inflammatory and anti-apoptotic role at the vascular tissue. In addition, previous studies have shown that NRF2 can also regulate the phenotypic switching of VSMCs through oxidative stress, but the specific changes and differences have not been studied at the cellular level. Also,there are studies believe that NRF2 is not necessarily completely protective of disease, disease-causing cells will also secrete NRF2 to play a self-protective role, resulting in the development of disease^23,64–66^.Therefore, it is necessary for us to accurately explore the role of NRF2 in VSMCs of AAA. VSMCs in AAA was first divided into NRF2+ group and NRF2-group according to its expression at sc-RNAseq level, in which NRF2-group accounted for the main proportion. Bioinformatic analysis results showed that NRF2+ group was in the upstream position of differentiation and expressed a large number of contractile and stress factors, which were mainly involved in cell contraction, oxidative stress and cell differentiation.While NRF2-group was in the downstream position of differentiation and expressed a large number of inflammatory factors and matrix remodeling proteins, which were mainly involved in inflammatory response and matrix remodeling. Subsequently, we further subdivided human aortic VSMCs into 7 subgroups, and NRF2 mainly had significant differences in contractile VSMCs between AAA and control groups, and ligand receptor analysis also showed that NRF2 could act on the classical contractile factor ACTA2. Therefore, we believe that AAA can regulate its phenotypic switching of VSMCs through NRF2 at scRNAseq level.

In the present study,VSMCs-specific NRF2 knockout significantly inhibited AAA formation, confirming a cell-specific role of NRF2 in AAA.Previous studies have reported that miRNA plays a significant regulatory role in AAA^67–69^.Through integrated scRNA-seq and transcriptomic profiling analyses,miRNA145 was identified as a downstream target trans-promoted by NRF2 and found to generate repressed effects of NRF2 on phenotypic switching,which being verified in vitro functional experiments. The miRNA145 coding gene, located on human chromosome 5, was thought to be co-transcribed in the same double-stranded transcript and has previously been reported to be large expressed in human heart and aortic VSMCs. Previous studies have reported that miRNA145 plays an important role in VSMCs lesions and phenotypic switching^70,71^.Moreover,it is believed that miRNA145 has become an effective way to treat cardiovascular diseases by regulating phenotypic switching.As early as 2009, a study published in nature showed that miRNA145 was a key factor regulating phenotype switching of VSMCs, which has high preventive and therapeutic value for acute myocardial infarction and hypertension^72^. In 2017, peiqiong zhou et al. believed that miRNA145 could effectively regulate VSMCs phenotypic switching, and vascular stents containing miRNA145 could be used to prevent vascular tissue regeneration^73^.In2019,research suggested that miRNA-145 can regulate VSMCs in the normal contractile phenotype, and inhibit the excessive proliferation and intimal hyperplasia to cope wih the restenosis in small-diameter vascular regeneration^74^.In 2021, Deborah D Chin et al. found that miR145 could rescue atherosclerotic protective systolic markers such as myocardium, α-SMA and calcitonin that synthesized in vitro, exerting its role in regulating phenotype transition and thus inhibiting the occurrence of atherosclerosis^75^. The most significantly altered pathway in VSMCs-specific NRF2 knockout mice is the MYOCD, which represents the phenotypic switching, so it is not difficult to guess the intermediate role played by miRNA145.

As a key protective factor for diseases, NRF2 has been widely used in clinic, such as aging prevention, tumor suppression, diabetes control and so on. In this study, the protective effect of NRF2 on AAA was confirmed by multiple aspects from bioinformatics analysis and animal experiments, which provided a new idea in the development of NRF2 targeted activation drugs for the treatment and prevention of AAA.Several limitations should be considered in this study. First, although a protective role of NRF2 in AAA has been confirmed in our study by using conditional knockout mice, the animal model is limited in its ability to mimic the extremely complex process of AAA formation in patients. In addition, although Tagln-Cre drivers have been commonly used to study VSMCs-specific expression, findings with this strain need to be interpreted with caution because its VSMC specificity is controversial.Second, the human AAA tissues obtained from advanced AAA lesions during open surgical repair provide limited insight into the earlier stages. Third, the interaction mechanism between NRF2 and miRNA145 was not deeply studied, although VSMCs phenotypic transition was an important mechanism of AAA development, the functional verification of miRNA145 in animal experiments of AAA was lacking in this study.More clinical and experiment researchs are warranted to confirm the benefit on AAA observed in this study.

## Nonstandard Abbreviations and Acronyms

AAA: Abdominal Aortic Aneurysm
Ang II: Angiotensin II
VSMCs: Vascular Smooth Muscle Cells
SMC: Smooth Muscle Cell
NRF2: Nuclear factor E2-related factor 2
ScRNA-seq: Single-cell RNA sequencing
EC: Endothelial Cells
IL-6: Interleukin-6
MMP2: Matrix Metallopeptidase 2
α-SMA/ACTA2: Actin Alpha 2, Smooth Muscle
CNN1: Calponin 1
SM22α: 22 KDa Actin-Binding Protein
MYOCD: Myocardium
WT: NRF2^flox/flox^
KO: NRF2^ΔVSMC^

## Highlights

1. We found that classical protective factors NRF2 prevent the development of AAA through the phenotypic switching of VSMCs.
2. We further verified the function of NRF2 at the cellular level by scRNA-seq combined with NRF2-knockout mice on VSMCs.
3. Mechanistically,NRF2 mediates phenotypic switching of VSMCs by modulating miRNA145/Myocd signal axis.

## Affiliations

Department of Emergency Medicine, Guangdong Provincial People’s Hospital (Guangdong Academy of Medical Sciences), Southern Medical University (X.X, X.H, F.S,G.C,Y.Z,X.L,Y.H,H.H,J.M,X.L). Medical Research Institute, Guangdong Provincial People’s Hospital (Guangdong Academy of Medical Sciences), Southern Medical University(X.H). School of Medicine, South China University of Technology (X.Z). Department of cardiovascular medicine department, Zhuhai People’s Hospital(LT.Z).

## Supplemental Material

Figures S1–S4

